# Route following by one-eyed ants suggests a revised model of normal route following

**DOI:** 10.1101/2020.08.21.261354

**Authors:** Joseph L. Woodgate, Craig Perl, Thomas S. Collett

**Affiliations:** Department of Biological and Experimental Psychology, School of Biological and Chemical Sciences, Queen Mary University of London, Mile End Road, London, E1 4NS, UK; Rothamsted Research, West Common, Harpenden, Hertfordshire, AL5 2JQ, UK; School of Life Sciences, University of Sussex, Brighton, BN19QG, UK; Department of Zoology: Functional Morphology, Stockholm University, Svante Arrhenius väg 18b, 106 91, Stockholm, Sweden

## Abstract

The prevailing account of visually controlled routes is that an ant learns views as it follows a route, while guided by other path-setting mechanisms. Once a set of route views is memorised, the insect follows the route by turning and moving forwards when the view on the retina matches a stored view. We have engineered a situation in which this account cannot suffice in order to discover whether there may be additional components to the performance of routes. One-eyed wood ants were trained to navigate a short route in the laboratory guided by a single black, vertical bar placed in the blinded visual field. Ants thus had to turn away from the route to see the bar. They often turned to look at or beyond the bar and then turned to face in the direction of the goal. Tests in which the bar was shifted to be more peripheral or more frontal than in training produced a corresponding change in the ants’ paths, demonstrating that they were guided by the bar, presumably obtaining information during scanning turns towards the bar. Examination of the endpoints of turns away from the bar suggest that ants use the bar for guidance by learning how large a turn-back is needed to face the goal. We suggest that the ants’ zigzag paths are an integral part of visually guided route following. In addition, on some runs in which ants did not take a direct path to the goal, they still turned to face and sometimes approach the goal for a short stretch. This off-route goal facing indicates that they store a vector from start to goal and use path integration to track their position relative to the endpoint of the vector.

## Introduction

Foraging ants, with their faculties intact, readily learn and follow visually guided routes between their nest and a foraging site (Collett et al. 1992, Wehner et al. 1996, Collett, M., 2010, Mangan and Webb 2012, Narendra et al. 2013). They can do so despite a large mismatch between their position on the route and their path integration (PI) state (Kohler and Wehner 2005, Graham and Cheng 2009, Narendra et al., 2013), implying that visual guidance does not require support from PI, even though the two guidance mechanisms are normally co-active (Collett, M., 2012; Wehner et al., 2016; Hoinville and Wehner 2018). Such experiments and modelling (Baddeley 2012) had suggested that visual route following involves ‘alignment image matching’ (AIM) (review: Zeil, 2012, Collett et al., 2013). In brief, ants that are guided initially by PI and by innate responses to obstacles and visual features that they encounter along the way, memorise routes by recording retinotopic views when facing along the route. Thereafter, they can follow the route by turning until they face in the direction that best matches a recorded view from the set of recorded views and then walking forward.

Since wood ants often use the same foraging path when going to and from their nest and on their outward route turn back to face along the path they have just followed, they can use what they have learnt on outward routes to help guide their return (Graham and Collett, 2006; Schwartz et al, 2020). A set of views may thus be associated with a particular motivational context so that an ant interrogates a different set on its approach to food than it does when homing (Harris et al., 2005; Wehner et al., 2005). A recently suggested alternative (Murray et al., 2020) is that ants activate both sets for guidance with one as attractive, mediating approach, and the other repulsive, causing avoidance, so improving the precision of route following. To account for the behaviour of wood ants from this perspective (Harris et al. 2005), the motivational context would determine which views are attractive and which repulsive.

Can one-eyed ants follow visually guided routes in the restricted surroundings of a laboratory lacking celestial compass information? In earlier experiments (Buehlmann et al. unpublished data), ants with one eye painted over were trained to find a drop of sucrose at the edge of a circular arena. They were released at the centre of the arena and learnt a straight route from there to a point on the circumference. This location was specified by a black vertical bar fixed to the white inner wall of a rotatable cylinder surrounding the arena. When the ants faced along the route, the bar was well outside the visual field of the ants’ seeing eye (e.g., Zollikofer et al., 1995). The cylinder and the position of the food were rotated together from trial to trial to ensure that the black bar was the principal visual cue and also to remove any reliance on a magnetic compass (Çamlitepe and Stradling, 1995). Ants with their left eye capped were approximately normal in route learning when the bar was on the right side of the route going from the centre to the periphery of the arena, but struggled when the shape was on the left side (Buehlmann et al. unpublished data). From the perspective of image alignment, this failure is not surprising. To the one-eyed ant, the view along the desired route is of an empty cylinder (Fig 1A) and the ant can fulfil this condition by picking from a wide swath of possible paths. One possible solution to the problem is to walk sideways while facing the cue (Fig 1B) – a strategy that we have not seen the ants adopt

**Figure 1.**
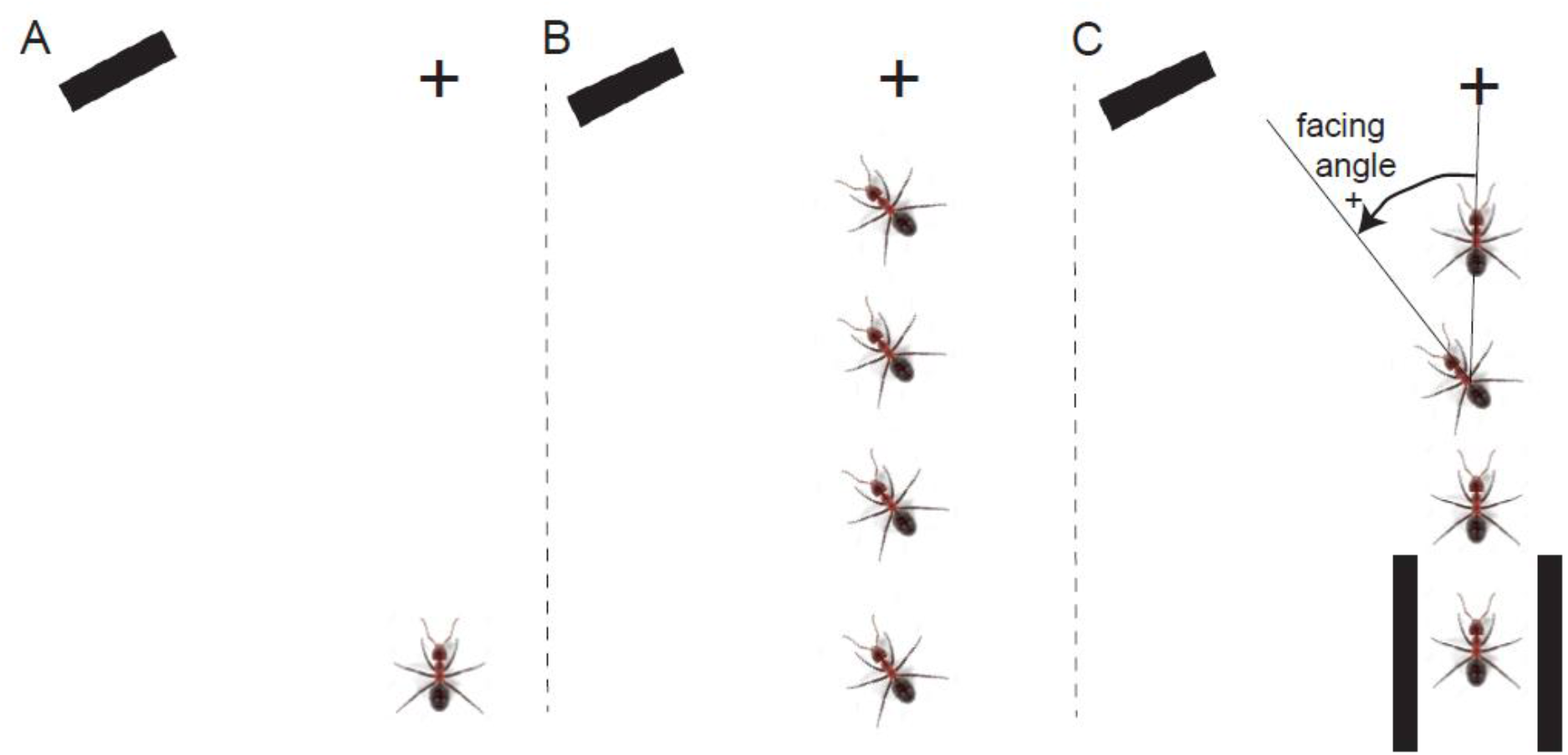
Schematic of problem and solution. **A**: Ant with left eye covered has difficulties in reaching its goal when the only directional aid is a black, vertical stripe outside the right eye’s visual field. **B**: Ant could, but does not approach, the goal when moving sideways. **C**: Ant is aided by mechanical guides forming a channel at the start of the route. It then turns to face towards and then away from the stripe and can reach the food (+).

In the present study, performed in 2017, we followed the same procedure except that we added extra information that seemed to help the ants learn a route. Ants were pointed in the right direction at the start of the route by two short parallel strips of wood aligned along the route (Fig. 1C). We examine the possibility that these guides help ants to develop a food vector (Collett et al., 1999; Bolek et al 2012) (i.e. a memory of the distance and direction from the start to the food); and that the ants’ routes are to some degree guided by PI, as they follow this vector. Tests with the vertical bar shifted further or closer to the route direction show that the ants also made use of the visual cue.

## Results

### Routes of monocular ants

Although ants were able to learn the route, the paths are much more variable than those of ants with normal vision guided by a similar cue (c.f. Woodgate et al. 2016). The directions of individual paths from the end of the channel to 15 cm from the food are shown plotted relative to the position of the food during training. If we select all the paths in the training trials before a test (Figure 2A, S1A), the bulk of paths are in a direction that lies within the expected quadrant (circular median: 2.85°; IQR: −22.35° to 11.69°). This value does not differ significantly from the direction of the food (binomial test, N: 131, P: 0.16; Zar, 1999). The details of the distribution are complex with a peak just to the right of the goal and another ca 20° to the left of the goal and subsidiary peaks close to the direction of the bar.

**Figure 2.**
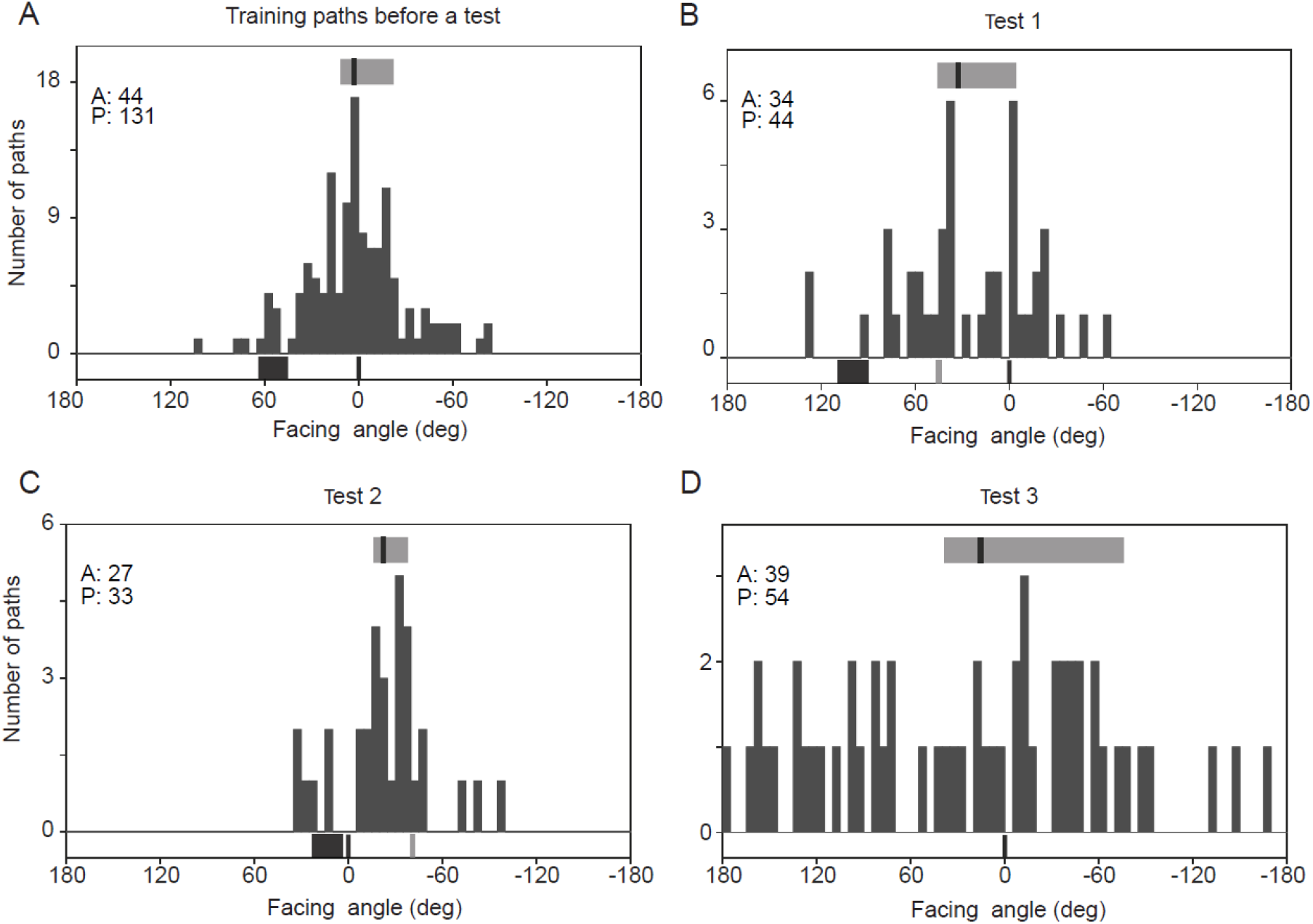
Path histograms. Path directions during training and three tests (see Methods). **A**: histogram showing the overall path directions taken by ants in training trials immediately preceding a test. Bin width is 5° in all histograms. Zero is the direction of the starting channel points from the centre of the arena (i.e., direction of food during training). In each histogram in this and other figures, A gives the number of ants and P the number of paths; the black line and grey rectangle above the histogram show the median and IQR of the distribution. The black rectangle below the histogram shows the width and position of the visual cue; the black line at 0° shows the food direction. **B**: path directions in tests with visual cue shifted left, relative to the starting channel. Black line at 0 shows the direction in which the starting channel was pointing; the grey line is the predicted goal direction if ants use the visual cue for guidance. **C**: path directions in tests with the visual cue shifted right, relative to the starting channel. **D**: path directions in the absence of the visual cue.

Tests in which trained ants were tested with the bar displaced to the left or to the right of its training position suggest that the ants’ direction of travel shifts with the displacement. In test 1, the right edge of the bar is about 90° from the direction in which the starting channel points (Figure 2 B, Figure S1 B). The number of paths is small (n= 44). One of two ‘major’ peaks (n=6) of the path directions (Figure 3 B) is 10° to the right of the expected goal relative to the shifted bar (bar-defined predicted heading: 45°). Another peak (n=6) is in the channel direction (channel-defined predicted heading: 0°). The circular median path direction (plotted relative to the channel-defined prediction; median 32.41 °; IQR −4.55 ° to 45.92°) differs significantly from the bar-defined goal (a), but does not differ from the direction of the channel (b) (binomial test, a: N=44, P: 0.010; b: N=44, P: 0.096). A comparison of path directions shows that the test 1 circular path median differed significantly from the median of training paths both when measured relative to the bar-defined goal direction (a) and relative to the channel direction (b) (Common median test, a: N=44,131, test statistic=10.11, P=0.001; b: N=44,131, test statistic=4.19, P=0.041).

**Figure 3.**
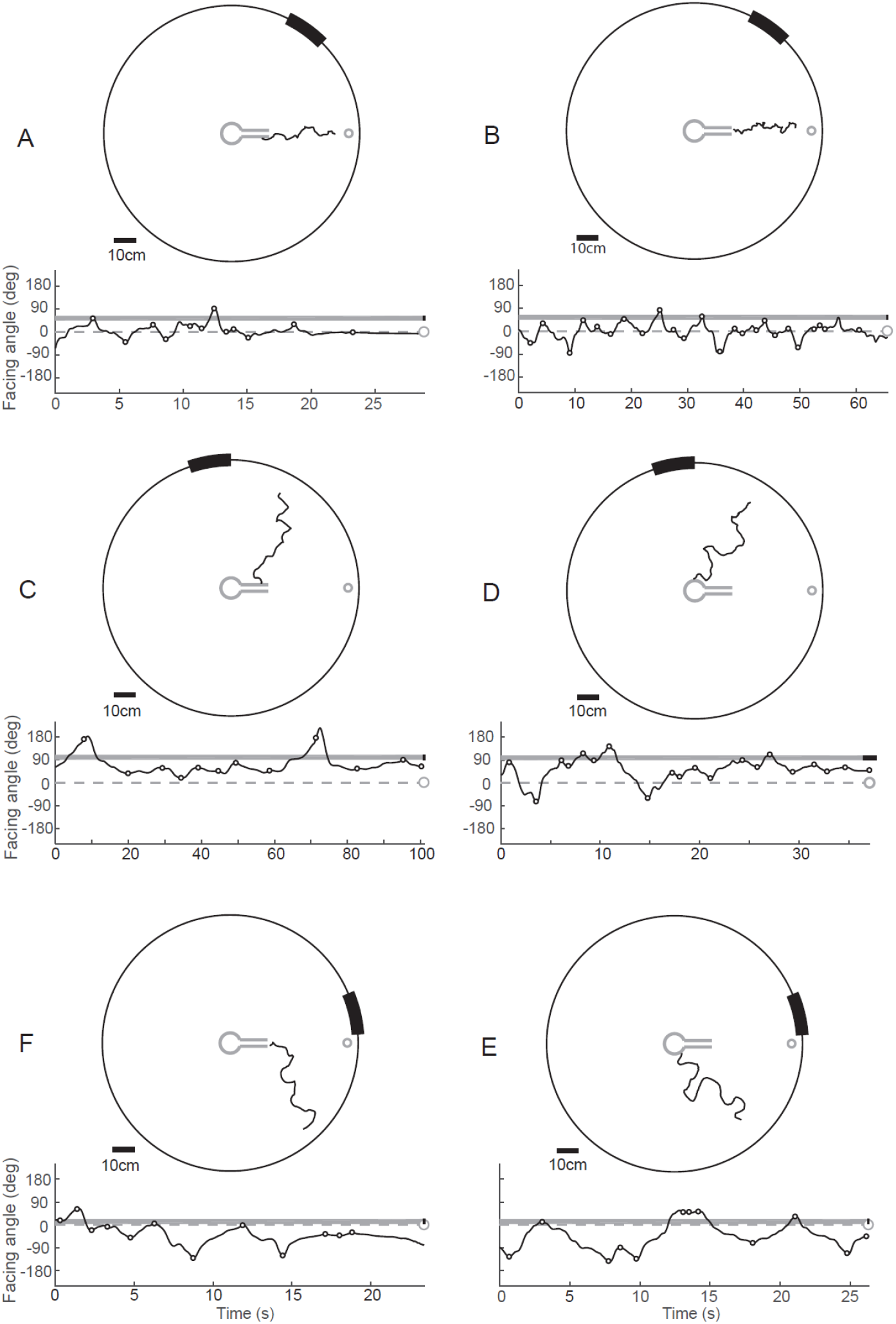
Individual examples of ants’ trajectories during training and tests. **A, B**: Training paths. Each panel shows the path of a different ant. Top: path of each ant, seen from above, as it moves from the enclosure and guiding channel in the centre of the arena towards the food position (0°) 55 cm away. Black bar on circumference of arena is the projection of the vertical bar that is fixed to the surrounding 180 cm diameter cylinder. Bottom: Ants’ facing angle relative to the food position (+ve angles counterclockwise) are plotted against time from start of the illustrated trajectory. Gray bar shows angular position of bar as measured from arena centre. Circles mark extracted turn extrema. **C, D**: Test with bar shifted more peripherally relative to the direction of the starting channel (no food present). Other details as in A. **E, F**: Test with bar shifted centrally relative to starting channel.

In a second test, the bar was moved to the right with its right edge almost in line with the channel. In this case, the paths shifted to the right of the training path (Figure 2 C, Figure S1 C). The peak of the distribution of path directions relative to the food lay between the training direction and the goal relative to the shifted bar (circular median −22.41°; IQR = −38.08 ° to −15.98 °, N:33). This median differs from both the bar-defined goal (a, bar-defined predicted heading: −41.2°) and from the channel direction (b, channel-defined predicted heading: 0°) (binomial test, a: N: 33, P: 0.0003 ; b: N: 33, P: 0.0003) The circular median of test 2 differs from the training path median, whether it is measured relative to the bar-defined goal direction (a) or the channel direction (b) (Common median test, a: N=33,131, test statistic=10.96, P=0.0009; b: N=33,131, test statistic=16.73, P<0.0001).

In the last test the bar was removed (Figure 2 D, Figure S1 D). The median direction is 15.54° (IQR = −75.87 ° to 38.74°, N 54) and does not differ significantly from the channel direction (binomial test, N: 54, P: 0.497). The dispersion of paths around the median was broader than in training (circular Watson U^2^ test, U 1.133; P<0.01; N: 54, 131). Nonetheless, the data are not uniformly distributed around the entire circle (Rayleigh test for uniformity: Z: 3.16, P: 0.042).

We conclude from these results that the ants’ path directions are influenced by the position of the bar, but that there are additional guidance mechanisms at play, such as a food vector.

### Individual paths

That the multiple peaks in the path direction data may be the consequences of following different navigational mechanisms is supported by the patterns seen in individual paths. Examples of test paths (Figure 3 C-F) indicate that ants can be guided by the bar, but also that guidance along the bar defined route can be accomplished with very few frontal views of the bar (Figure 3C). Training routes with few frontal views of the bar (Figure 3A) could mean that ants, guided idiothetically, without compass information, follow their starting directions to the food (e.g. Lent et al., 2009). Ants may also be guided idiothetically by PI, as in *Drosophila* returning to a drop of sucrose that they have sampled (Kim and Dickinson, 2017).

Suggestive evidence for a food vector monitored through PI comes from examples of ants that leave the direct route to the goal and turn and travel towards the food when they are some distance from the trained path. Deviations from the trained path happens when ants head towards the bar (Figure 4 A-C, G-J) or travel well to the right of the direct route from the centre to the food (Figure 4 D-F). These deviant paths can be interrupted by the ant turning towards the goal and moving a short (e.g. Figure 4, F, G) or a longer distance (e.g. Figure 4 E, I) towards it. When the distance towards the goal is longer, and is not just a brief interruption to a path elsewhere (Fig. 4 D, E, H), facing the goal tends to occur at the peaks and troughs of a zigzag approach, as it does in binocular ants (Lent et al., 2013).

**Figure 4.**
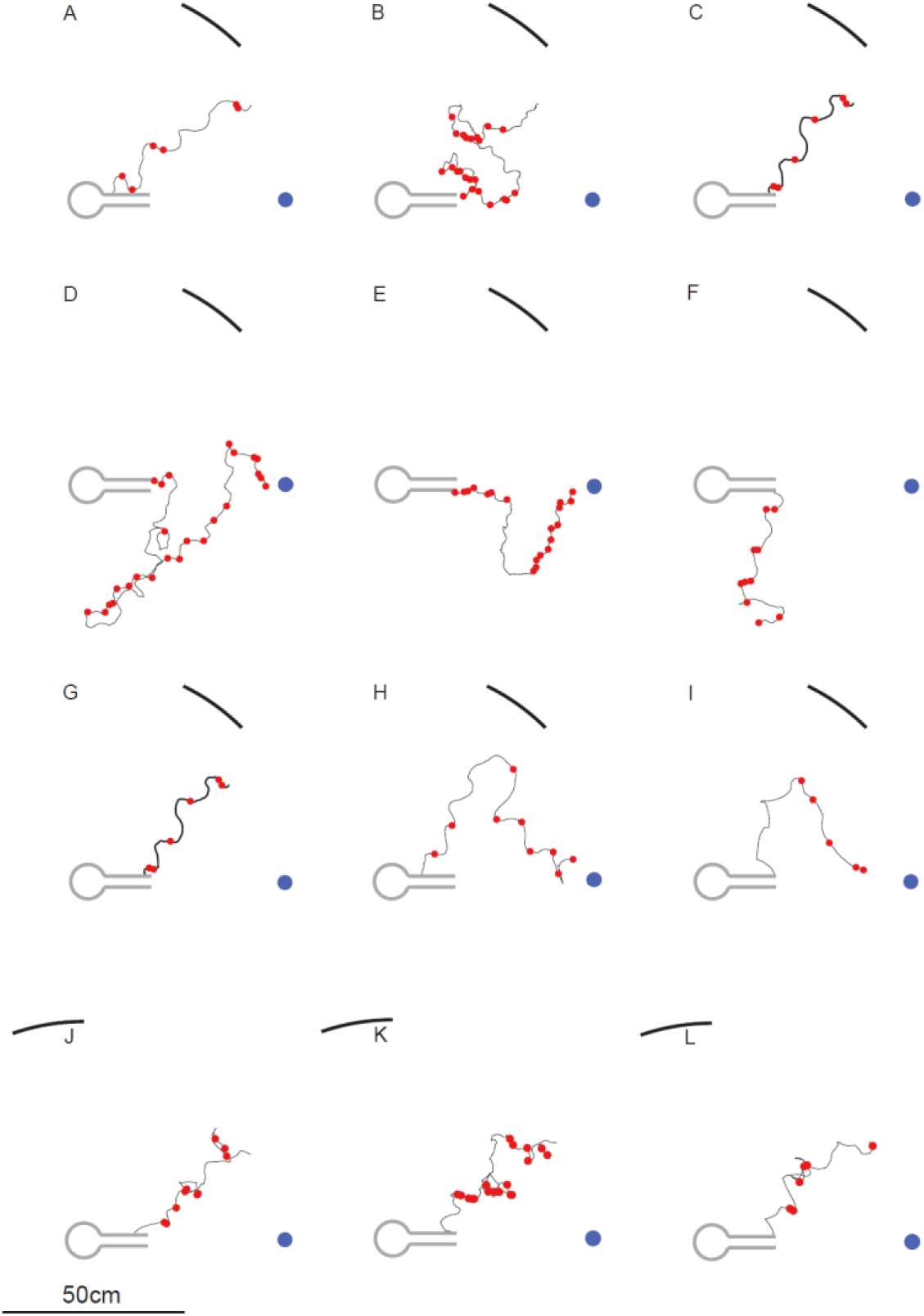
Suggestive evidence that ants can be guided by path integration to the food position. Example paths of ants that deviated far from the direct path to the bar defined goal. Points or path segments in which the ants face the goal within ±10° are marked with red dots. Panels **D, E, H** and **I** show ants that stray far from the direct route between the start and the food, but later make move directly toward the goal, suggestive of the action of a food vector monitored by PI. Other panels show shorter periods of movement in the goal direction and illustrate the range of circumstances under which goal facing occurs. Panels **J, K** and **L** show similar goal-facing in tests in which the position of the visual cue has changed, demonstrating that this behaviour cannot be entirely visually mediated.

The examples mentioned so far come from training trials with a drop of sucrose at the goal. Could the ants obtain guidance cues from the sucrose, itself? During experiments over many years in this set up, ants never seem to notice the food until they have almost reached it. These concerns do not arise during tests in which food is always absent. Ants continue to face the channel-defined goal when approaching the bar in test 1 (Figure 4 J-L).

Facing and moving towards the goal can most easily be explained by supposing that ants learn a vector that goes from the centre of the arena to the goal and rely on PI to monitor their progress along the vector and to generate a home vector, whatever path they take (cf. Fernandes et al., 2015). The examples in Figure 4 also show how readily ants switch between following PI and controlling their path with a visual cue.

### Facing directions at the end points of turns

The distributions of path directions (Figure 2) indicate that the bar has a major influence on the ants’ direction of travel. More detail comes from examining separately the endpoints of the left and right turns that ants make during their paths. We extracted two kinds of turn endpoints from the data: i) the maxima and minima in plots of the ants’ facing angles (e.g. Figure 3B); ii) turns that end in a plateau, where the ant keeps its facing direction steady for at least 0.2s (e.g. Figure 3A) and combined the two kinds of endpoints in all the histograms below (Figure 5).

**Figure 5.**
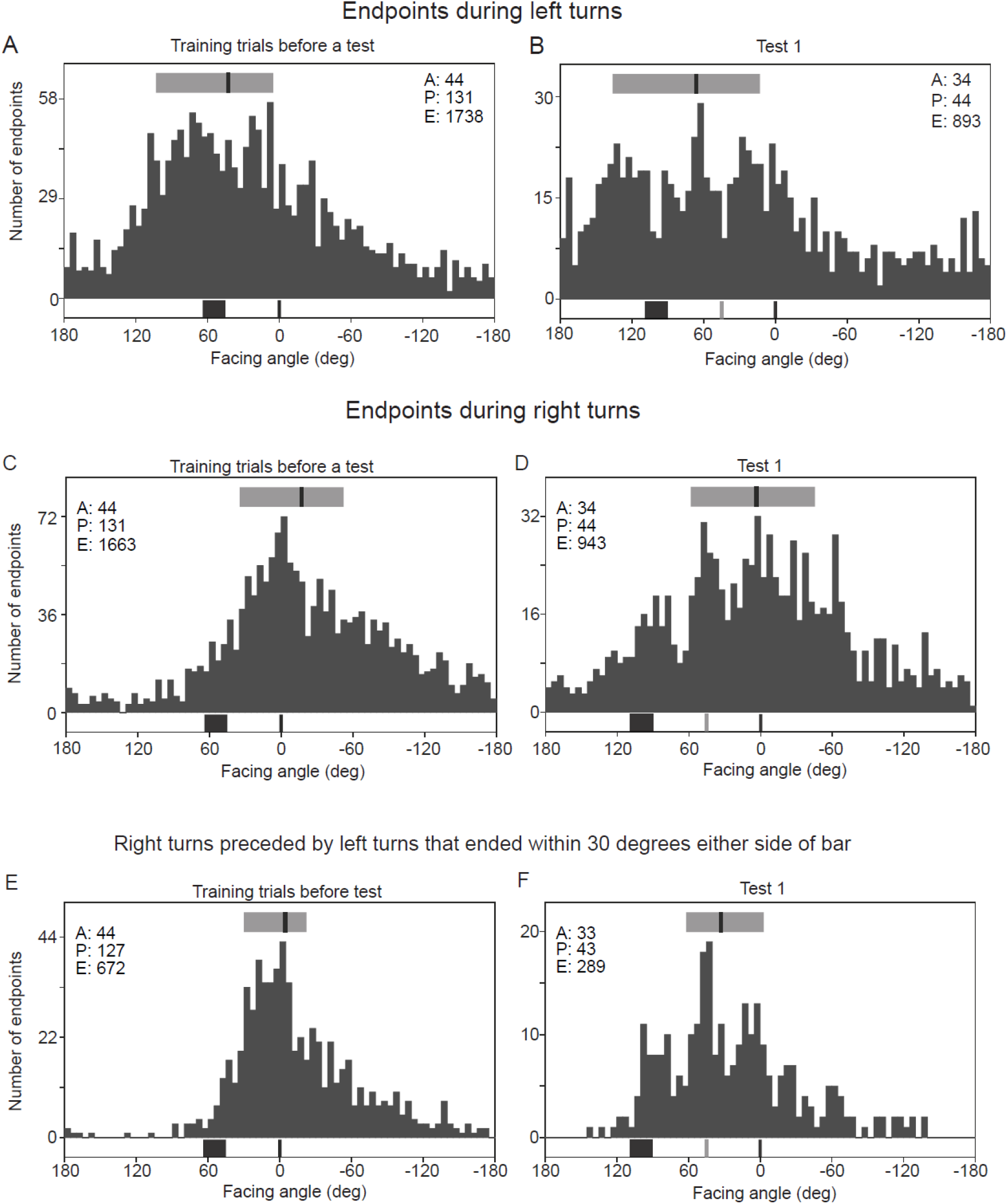
Histograms of endpoints of left and right turns. **A**: All endpoints of left turns during pre-test training trials. **B**: Endpoints of left turns during Test 1 trials, with visual cue shifted left relative to the starting channel. **C, D**: Endpoints of right turns in pre-test training and Test 1, respectively. **E, F**: Endpoints of selected right turns that began ±30° from the bar edges. In all panels, E denotes number of endpoints. Other details as in Figure 2

Despite the wide spread of the distributions, the pattern of peaks is informative. In the pre-test training trials, the distributions of left turn endpoints have a clear peak associated with the food which is likely to reflect turns generated through PI (Figure 5A). A second peak consists of left turns that end close to the bar. The endpoints of right turns during training are in the food direction and these end points could either be generated through PI or reflect a right turn of a remembered size after facing the bar (Figure 5C).

In test 1, the endpoints of left turns are shifted to the left in comparison with training, but the details of the pattern are too noisy to be informative (Figure 5B). The distribution of endpoints of right turns has two peaks, similar to the pattern of path directions during test 1 (Figure 5D, c.f. Figure 2B). One peak is in the bar-specified goal direction and a second is in the channel-specified direction. The peak in the bar-specified goal direction is enhanced, when we select just those right turns that occur after viewing the bar (Figure 5F). To do so, we included only endpoints that followed turns ending within 30° either side of the edges of the bar. The comparison of this plot with the plot of all right turns during test 1 provides our best evidence that ants do learn the amplitude of the turn that they must make to face in the goal direction after viewing the bar.

Since we cannot suppose that turn endpoints are independent of each other, we have not attempted statistical analysis. The same pattern is seen when we look at just the median endpoint from each ant path (Figure S2). The small sample number and variability of the data means that the only useful comparison to be made is whether the circular median directions of test 1 (median, relative to the bar-defined predicted goal = −18.2° IQR = −31.9° to 14.2°, 43 paths) differ from that of test 2 (median, relative to the bar-defined predicted goal = 0.8°; IQR = −13.7° to 13.5°; 32 paths). We selected all the median endpoints from ants that undertook both tests and aligned the distributions on the bar-defined goals. A common median test demonstrated that the two distributions differed significantly (Common median test, N: 19, 19, test statistic: 5.157, P: 0.023, Fisher, 1995). We conclude that the endpoints of right turns cannot be driven by the position of the visual cue alone but must also be influenced by the mismatch between the bar and channel positions.

Right turn endpoints during test 2 peak at a position that falls between the bar-specified and channel-specified directions. This compromise may be another example of ants averaging directions of the outputs of independent navigational strategies (visual guidance and PI or idiothetic guidance) when the directions of the two outputs are relatively close (Collett, 2012; Wehner et al., 2016; Hoinville and Wehner, 2018).

### Binocular ants

We have re-analysed an earlier study on binocular ants to see whether the findings on one-eyed ants apply to normal route following. Wood ants (Woodgate et al 2016) were trained in the same apparatus to approach a goal with the direction set by a single rectangle placed to the left of the goal direction and tested with wider rectangles. The major result of various tests was that ants seemed to learn their route relative to the centre of mass of the shape. We now ask whether turn endpoints also suggest that ants learn the goal direction relative to the centre of mass of the training shape and wider test shapes.

We extracted the endpoints of selected turns during training paths preceding a test and during tests (Figure 6). The selected endpoints of left turns were those that began within ±20° or ±30° of the predicted goal direction from the midpoint of the width of the training rectangle and wider test rectangles. Selected right turns began within ±20° or ±30° of the midpoint of these rectangles. As with monocular ants, the distribution of endpoints is complex and multi peaked. Major peaks of left turns occur a little to the right of the midpoint of the training and wider rectangles and peaks of right turns are a little to the left of the goal direction relative to the midpoint of the rectangle. Given the complexity of the data, we cannot say more than the data are consistent with the importance of the midpoint of the rectangles as one factor influencing the pattern of endpoints and the turns that the ants make.

**Figure 6.**
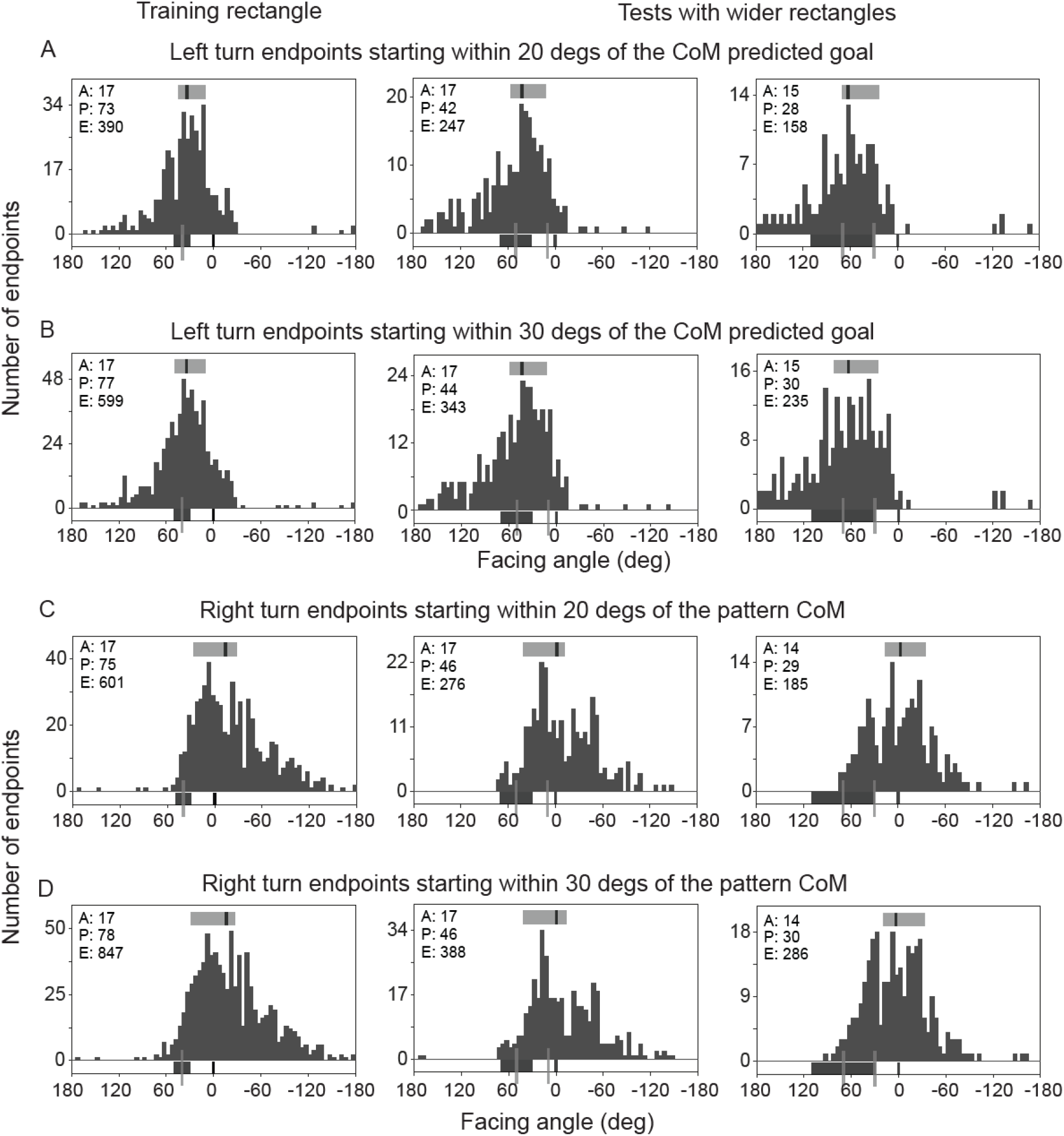
Histograms of endpoints of binocular wood ants during selected left and right turns. Endpoints of turns in pre-test training trials, tests 1 and tests 2, in which rectangles were increasingly wide. **A**. Left turns with their starting points within ±20° of the predicted facing angle of the goal relative to the centre of mass (CoM) of each rectangle. Black lines below the histograms at 0° show the predicted goal direction during training. This direction is also the predicted direction were ants to match the right edge of the rectangle to its position in training. Grey lines below the histograms mark the CoM of the visual cue and the predicted goal direction, were ants to match the CoM of the rectangle to its position in training. Other details as in Figures 2 and 5. **B**. Left turns with starting points within ±30° of the predicted facing angle of the goal relative to the CoM of each rectangle. **C**. Right turns with starting points within ±20° of the CoM of each rectangle. **D**. Right turns with starting points within ±30° of the CoM of each rectangle.

An earlier study of ants approaching visual features aligned with the route found that facing along the route direction tended to occur at the apices of the ants’ zigzag path (Lent et al. 2013). Examples of paths from (Woodgate et al 2016) show that moments when the ants face the goal head on coincide closely with the peaks and troughs of the ants’ paths (Figure 7). The coincidences occur even when the zigzags are irregular.

**Figure 7.**
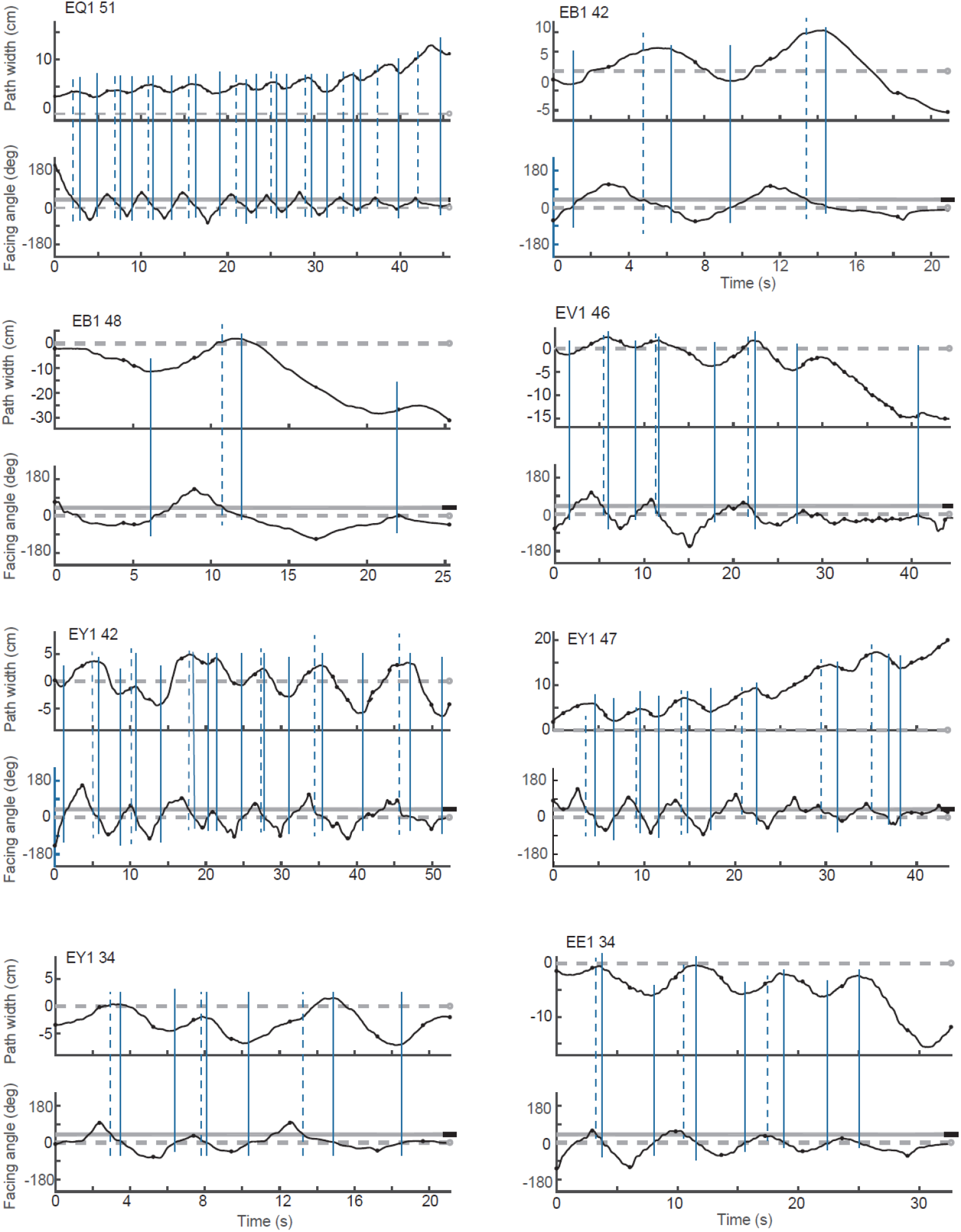
Examples of zigzag paths in binocular wood ants. Each panel gives one path. Top trace: ant’s position along its path plotted against time. Horizontal grey dashed line shows the direct route from the starting point to the food. Bottom trace shows ants’ facing angle relative to food. Horizontal dashed grey line shows when facing angle is directly towards the food; grey solid line represents the angular position of the bar as measured from the centre of the arena. Continuous vertical blue lines highlight points at which the ant is facing directly toward the food. These lines tend to intersect the peaks and troughs of the ant’s Y position irrespective of the spatial or temporal regularity of the zigzag. Dotted blue vertical lines highlight points at which the ants faced toward the bar during turns in the direction of the food and show the variability of the interval between the ant facing the bar and facing the food.

## Discussion

Alignment image matching, the prevailing account of visually guided navigation in insects, suggests that an animal will turn until its current view matches a previously learned view and then move in that direction (Baddeley et al., 2012). In our study, that process could not operate because one eye was painted over and the only visual cue was on the blind side of the ant. Consequently, when the ant faced in the correct direction, it saw no more than the white arena wall. Ants still learnt routes to a feeder and, if the visual cue was shifted in tests, the ant’s paths were deflected in the appropriate direction, demonstrating that the ants’ movements were guided by the visual cue.

A wood ant’s path tends to oscillate with the desired goal direction reached at the peaks and troughs of the oscillations or zigzags (Lent et al. 2013). This is a common pattern in insect locomotion. It is of clear benefit to insects that move in an oscillating path to follow an odour trail (Hangartner, 1968; Cardé and Willis, 2008; Wllis et al, 2013; Namiki and Kanzaki, 2016). When one-eyed ants zigzagged in the current experiments, they would see the bar during turns to the left, giving them an opportunity to acquire information to guide their subsequent routes. We suggest that, during route learning, the right turns after turning to face the visual cue were guided by path integration so that turns ended with ants facing towards the goal. This turn size can then be learned and later applied when following an acquired route. By this means, visual scanning can support AIM and make it possible to navigate in surroundings with sparse visual information.

The coincidence of facing the along the goal direction with the apices of the zigzag, even when the zigzag is irregular suggests that turns in the goal direction help induce a reversal in the ants’ direction of rotation. Because the ants often sustain their direction towards the goal, the influence of goal facing on controlling the direction of rotation seems to be more of a slowly acting nudge than a kick.

One puzzle that the data present is that although ants seem able to reach the goal by following a food vector without needing the bar, they mostly did not travel in the food direction in tests with the bar absent (Figure 2D). A possible explanation comes from previous work on the expression of home and goal vectors that suggests that the vectors are not fully expressed if the surroundings in which they are tested differs markedly from the ant’s accustomed surroundings (Cheung et al., 2012). The bar may thus provide a contextual, scene-setting cue as well as a cue for guidance. Another possibility is the lack of optic flow that the bar normally provides.

While AIM as originally conceived is likely to be mediated by the mushroom body (Ardin et al, 2016, Buehlmann et al., 2020; Kamhi et al., 2020) that signals an attractive or aversive direction of travel (Aso et al., 2012), controlled turns to or from specific visual features are probably mediated by visuomotor processing in the central complex (CX) (Collett and Collett, 2018). Visual learning of pattern orientation and elevation was some time ago shown to involve ring cells in the ellipsoid body (EB) and cells in the fan-shaped body (Liu et al., 2006; Pan et al., 2009). The experimental technique of these experiments prevented testing whether the azimuth of a visual stimulus is also memorised within these structures.

Current research on the *Drosophila* CX has resolved this question in wonderful detail. First, it was found that a fly’s heading direction is correlated with the position of a single ‘bump’ of electrical activity within the doughnut shaped EB, and that the position of the bump can be locked to a compass direction or to a visual stimulus (Seelig and Jayaraman, 2015; Turner-Evans, D. 2017; Green et al., 2017, Giraldo et al., 2018).

More recent imaging studies and recordings from ring and EP-G neurons in the EB have uncovered the likely mechanism that underlies how the bump of activity within the EB becomes linked to the direction of a visual cue (Fisher et al., 2019; Kim et al., 2019). Individual ring cells in the EB respond to visual cues in a specific retinal position. When a ring cell is visually activated, it inhibits E-PG cells at all positions within the EB, apart from where the ring cell is itself visually activated. This system is plastic such that E-PG cells responsive at a specific heading become linked to a visual cue in that direction after that cue has remained in a fixed direction for a short time. This arrangement allows a facing direction within a visual scene to be rapidly associated with a particular compass direction, just as is required for a visually controlled route learnt through path integration.

The turn amplitudes that are likely to be mediated by known CX circuitry are of several kinds. Turns that place a visual stimulus in its expected position, like turns towards and away from the bar, can be understood within the framework of the previous paragraph (Fisher et al., 2019; Kim et al., 2019). Turns in the direction of the home vector are also likely to involve the CX, in this case, interactions between the fan-shaped body, the protocerebral bridge and the EB (Neuser et al. 2008; Stone et al., 2017; Honkanen et al 2019, Le Moël et al., 2019). Lastly, the lateral accessory lobes that contain neurons that mediate the turn of a zigzag path (Namiki et al. 2016, Steinbeck et al, 2020) may be responsible for the correlated phasing of goal facing and the peaks and troughs of zigzags (Figure 7).

## Methods

### Ants

Experiments were performed on wood ants from laboratory-maintained colonies collected from Broadstone Warren, East Sussex, UK. The colonies were kept under a 12h:12h light-dark cycle and the colony was sprayed with water daily. Water and sucrose dispensers were always available except during experiments when the colony had limited access to sucrose to encourage enthusiastic foraging. Frozen crickets were supplied several times a week.

Before ants were trained their left eye was painted with enamel paint and the integrity of the paint cover checked daily under a binocular microscope. The cover appeared stable and it was very rare to have to repaint.

#### Experimental set-up

The basic experimental procedures followed those described previously (Lent et al., 2013). Individually marked wood ants were trained to go from the centre of a circular platform (radius 60cm) towards a drop of sucrose on a microscope slide placed 55cm from its centre. The slide was positioned relative to a single black vertical bar (50cm wide and 85cm high) that was cut from black cotton sheeting and fixed to white netting on the white inner wall of a rotatable cylinder (diameter 3m, height 1.8m). Seen from the centre of the arena, the right edge of the bar was 45° to the left of the direction the feeder.

To provide idiothetic cues, a pair of guide sticks (3mm square cross-section) with a 24mm path between them led from a ring (5cm radius and 25mm high) surrounding the centre of the arena. In initial training trials, the sticks led most of the way to the food, their length was gradually shortened to 13cm. 13cm and 10cm sticks were used for tests. See Fig. 3 for a top view of the arena the arrangement of sticks, the sucrose and the radial position of the bar.

#### Experimental procedure

Ants were given about 25 trials of group training over about 3 days before being trained individually. On each trial, individually marked ants from a cohort of *ca.* 25, were taken from the nest and separated into groups of between 6 and 8 individuals. One group at a time was placed in the ring, sometimes overlapping with stragglers from a previous group. After the ants had reached the feeder and had started to drink, they were placed in a box with sucrose and drank to completion before being returned to the nest.

For individual training, ants were put singly into a 6.5cm diameter, cylindrical holding chamber that lay within the ring. The wall of the holding chamber could be lowered remotely from outside the cylinder to be flush with the arena floor. Once the wall was lowered the ant was free to leave the ring. When the ant had reached the food reward and started to feed, or reached the edge of the arena, the experimenter raised the wall of the holding chamber, entered the cylinder, transferred the ant to a feeding box and placed the next ant in the holding chamber. After the cohort of ants had completed a training trial, the guide sticks, the black bar and the slide with food were rotated to a new position to avoid the ants learning other cues for guidance.

Once individual training began, we recorded the ants’ individual paths, starting just before each ant was released from the holding chamber. Each ant’s movements were recorded using a tracking video camera (Trackit, SciTrackS GmbH), which gave as output the ant’s position on the arena and the orientation of its body axis every 20ms. Each ant was released from the holding chamber and its path recorded until it reached the feeder or the edge of the platform. Once it had started to feed, or if it was lost, it was placed in a feeding box.

After individuals had performed 30 training trials, three different tests were introduced with a varying number of intervening training trials. In test 1, the right edge of the bar was shifted to 90° to the left of the direction in which the starting channel was pointing. In a second test the bar was shifted in the opposite direction, ca 5° to the left of the channel direction. In the third test the bar was removed. No sucrose reward was present during tests. Ants were removed from the arena once they reached the edge of the platform and allowed to feed to satiety before being returned to the nest.

#### Data analysis

We examined two features of the ants’ trajectories: 1. The direction of an ant’s path; 2. The ant’s facing directions at the extrema of its turns to the left and the right, measured relative to the position of the food (see Figure 1C).

##### 1. Path direction

The direction of the ants’ paths was analysed from when an ant left the guide channels, beginning ca. 18cm from the centre of the arena, or sooner if ants climbed over the guide strips, until they reached a radial distance of 45cm from the centre. The direction over this path-segment was estimated in two stages. First, the path (consisting of x-y coordinates recorded at 50fps) was divided into successive 1.5cm segments, and orthogonal distance regression (Gene et al., 1996) was used to calculate the angle of a best fit line through each segment, minimising the error in both x and y dimensions (custom written MATLAB script based an algorithm devised by Eitan, T. [https://stackoverflow.com/questions/12250422/orthogonal-distance-regression-in-matlab]). The overall direction of the path was then taken to be the vector sum of the directions of all path segments until the ant’s path took it 45cm from the centre of the arena.

##### 2. The amplitude of scans

Plots of facing angles over time tend to have peaks and troughs. The facing angle at a peak or a trough was defined as the most extreme angle to the left or right before a change in the direction of rotation. These extrema were identified using the following procedure: first, to reduce noise in the data, the ant’s facing directions throughout each track were smoothed by taking the circular mean facing direction across a 50-frame moving window. Points at which the angular change between consecutive frames reversed direction gave the rough location of inflection points in the track. To pick out the facing direction just prior to a reversal of direction, we used the unsmoothed angle data to find the interval over which the facing angle was more extreme than the facing angle at the inflection point. The mid-point of this interval was taken as the facing angle at the peak or trough.

The algorithm used to extract extrema does not capture all the turn-endpoints that can be seen by eye (e.g. Figure 3), because some turns were too small for the inflection point to show up in the smoothed angle data. We saw no indication of false identification of extrema.

Some turns end in a clear plateau rather than a turn in the opposite direction. When this was the case, the ant’s facing angles at these plateaux were added to the extrema data to give what we term turn endpoints. Plateaux were identified through the fulfilment of two criteria: (1) an interval during a track that lasted at least 0.2s, in which the maximum change in facing angles was less than 3°; (2) that the slope of the regression of facing angle against time of a candidate plateau should be less than 1°s^-1^. The facing angle of an identified plateau was then defined as the circular mean of all facing angles in the interval.

#### Exclusion of inconsistent ants

Some ants were very erratic. Paths of these ants are included in Figure S1 but excluded in further analysis. To remove these ants, we calculated the overall direction of travel of each path from every ant and calculated the resultant vector length of the path directions of each ant across the entire training period. Ants whose resultant vector length >0.4 took similar directions, relative to the visual and idiothetic cues, in each trial, despite the rotation of the experimental arena. Those with a vector length <0.4 could not cope with rotation; 5 such ants were removed from the training and test dataset. One of these 5 had taken part in tests 1 and 2, and none had participated in test 3.

### Descriptive statistics

At the top of each histogram, we show the circular median of the distribution (black line; Otieno and Anderson-Cook, 2003) and the interquartile range (IQR; grey bar; defined for circular data, using the likelihood-based arc method, as the smallest arc of the circle containing 50% of all datapoints, Fisher and Hall, 1989). Rayleigh’s tests for non-uniformity confirmed that the ants’ path directions and endpoints in training and tests 1 and 2 were not uniformly distributed but showed a tendency to head in one direction (all P<0.0001).

## Supplementary figures

**Figure S1.**
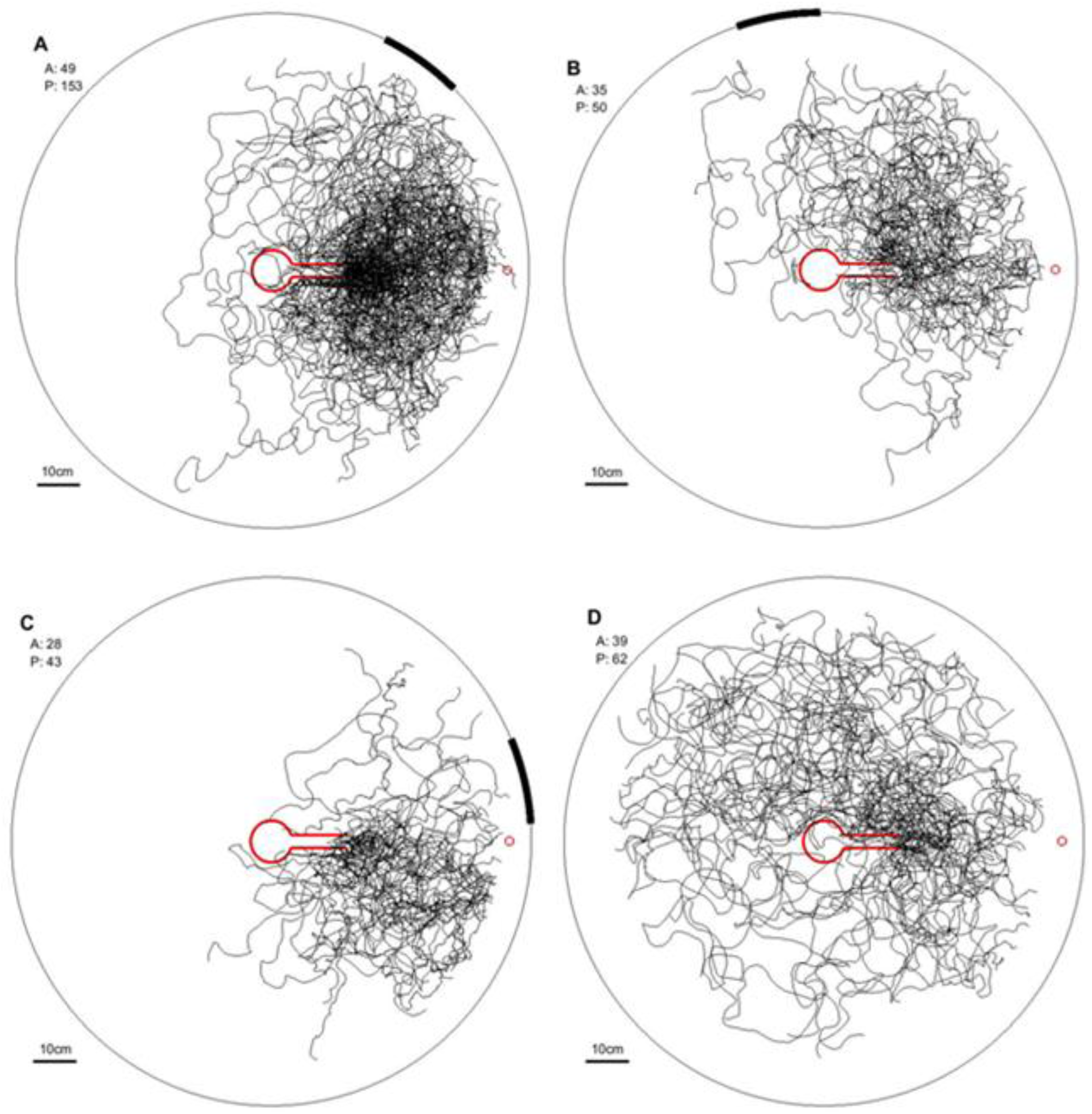
Superimposition of ants’ paths during training conditions and tests. All paths in training trials just before a test and in tests 1-3 are shown superimposed without any filtering for consistency (See ‘Exclusion of inconsistent ants’ in Methods). **A.** paths recorded in training trials **B.** Paths in test 1 with bar shifted away from food. **C.** Paths in test 2 with bar shifted towards the food. **D.** Paths in test 3 with the bar removed. In each panel, A gives the number of ants and P the number of paths.

**Figure S2.**
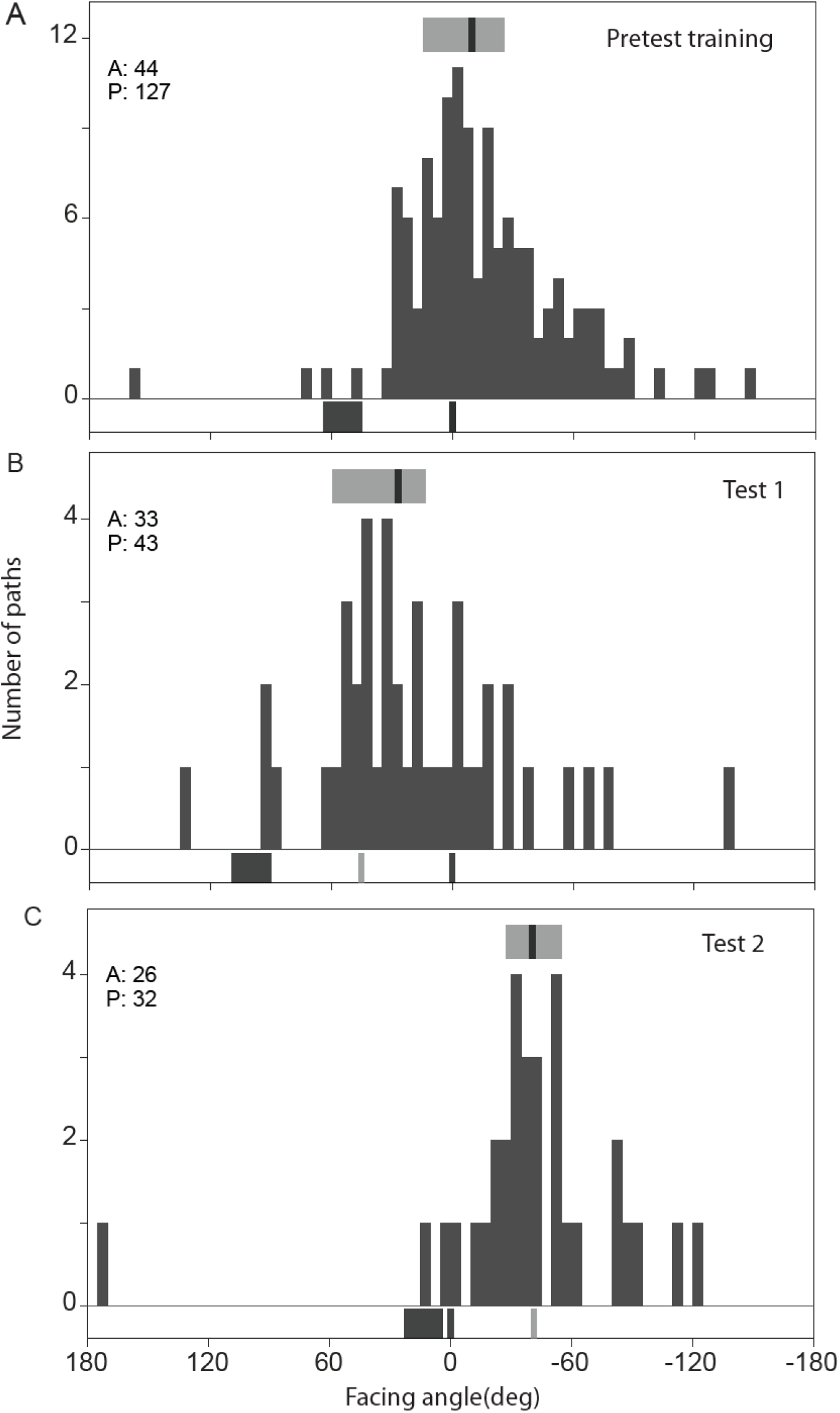
Replot of endpoints of selected right turns as in Figure 5. **A**: Histogram showing, for every pre-test training path, the median of the endpoints of selected right turns within the path. Within each path, selected turns began ±30° from the bar edges. Each data point represents one path. The pattern of peaks resembles that of Figure 5 **B**: Median endpoints of selected turns in test 1 trials, with the visual cue shifted left, relative to the starting channel. **C**: Median endpoints of selected turns in test 2 trials, with the visual cue shifted right, relative to the starting channel.

